# Telomere dysfunction represses HNF4α leading to impaired hepatocyte development and function

**DOI:** 10.1101/619601

**Authors:** Evandro L. Niero, Wilson C. Fok, Alexandre T. Vessoni, Kirsten A. Brenner, Luis F.Z. Batista

## Abstract

Telomere attrition is a risk factor for end-stage liver disease. Due to a lack of adequate models and intrinsic difficulties in studying telomerase in physiologically relevant cells, the molecular mechanisms responsible for liver disease in patients with telomere syndromes remain elusive. To circumvent that, we used genome editing to generate isogenic human embryonic stem cell lines (hESCs) harboring a clinically relevant mutation in telomerase (DKC1_A353V) and subjected them to an *in vitro*, stage-specific hepatocyte differentiation protocol, that resembles hepatocyte development *in vivo*. Our results show that while telomerase is highly expressed in hESCs, it is quickly silenced, due to *TERT* down-regulation, after endoderm differentiation, and completely absent in *in vitro* derived hepatocytes, similarly to what is observed in primary hepatocytes. While endoderm derivation is not impacted by telomere shortening, progressive telomere dysfunction impaired hepatic endoderm formation. Consequently, hepatocyte-derivation, as measured by expression of specific markers, as well by albumin expression and secretion, is severely compromised in telomerase mutant cells with short telomeres. Interestingly, this phenotype was not caused by cell death induction or senescence. Rather, telomere shortening induces down regulation of the human hepatocyte nuclear factor 4α (*HNF4α*), in a p53 dependent manner. Telomerase reactivation, as well as p53 silencing, rescued hepatocyte formation in telomerase mutants. Likewise, conditional expression of *HNF4α*, even in cells that retained short telomeres, accrued DNA damage, and p53 stabilization, successfully restored hepatocyte formation from hESCS.

**Conclusions:** Combined, our data shows that telomere dysfunction acts a major regulator of *HNF4α* during hepatocyte development and function, pointing to a potential novel target for the clinical management of liver disease in telomere-syndrome patients.

## INTRODUCTION

Telomeres are repetitive DNA sequences that prevent degradation and fusion of chromosomal ends. As the DNA replication machinery is unable to fully replicate DNA termini, telomeres become progressively shorter with consecutive cellular divisions, eventually reaching a critical stage where telomere dysfunction induces senescence or cell death^1^. Evidence collected during the past decade has established impaired telomere maintenance as a causative effect in a number of different conditions, ranging from bone marrow failure to pulmonary fibrosis and liver disease^2^. Interestingly, telomere shortening affects different tissues at different ages, and telomere length measurement in hospital settings has recently been proposed as a diagnostic tool in targeted indications, with the potential to inform treatment decisions and influence morbidity^3^.

The active telomerase complex is composed of *TERT* (reverse transcriptase component), *TERC* (the telomerase RNA component), and dyskerin (*DKC1*), necessary for *TERC* stabilization^4^. While mutations in these genes were initially found in children suffering from the bone marrow failure syndrome dyskeratosis congenita (DC), adult-onset phenotypes, such as pulmonary fibrosis and liver disease now represent the most common presentation in patients harboring mutations in telomerase. Interestingly, patients with liver disease usually come to clinical attention at an earlier age and with longer telomeres when compared to patients with pulmonary fibrosis^3^, indicating that liver cells might be more susceptible to telomere dysfunction than the lung epithelia. Liver abnormalities have recently been shown to be common in patients with telomere syndromes, with liver disease carrying important implications for morbidity and mortality in these patients^5^. Accordingly, telomerase deficient mice which underwent ablation of the liver showed impaired hepatocyte regeneration and accelerated development of liver cirrhosis after chronic liver injury^6^. In addition, it has recently been shown that a small population of *TERT* positive hepatocytes are responsible for hepatocyte renewal and maintenance of liver mass, both in homeostasis and after critical injury^7^. While these results highlight the importance of telomerase for hepatocyte regeneration and liver disease, the chain of events linking telomere shortening to hepatocyte and liver failure in humans remains elusive. In fact, while genetic mutations in telomerase have been associated with different liver diseases, ranging from non-alcoholic fatty liver disease and non-alcoholic steatohepatitis to fibrosis and cirrhosis, the mechanisms behind liver tissue reorganization and failure in settings of dysfunctional telomeres have not yet been elucidated^8, 9^. A major hurdle to address this question has been a lack of viable models to specifically analyze the impact of exacerbated telomere shortening in human hepatocytes. To overcome this limitation, we used the targeted differentiation of human embryonic stem cells (hESCs) into mature, hepatocyte-like cells. We used wild-type (H1 hESC line) and CRISPR/cas9 isogenic engineered hESCs harboring a clinically relevant mutation in telomerase (DKC1_A353V mutants created by introduction of a 1058C>T point mutation in the *DKC1* gene^10^). The engineered cells are free of detectable chromosomal abnormalities by G-band analysis and are pluripotent, but have reduced telomerase activity and progressive telomere shortening due to low levels of *TERC* expression (Supplemental Figure 1A-G).

## METHODS

### Cell Culture

H1 (WA01) hESCs were acquired from the WiCell Research Institute (Madison, WI), following all institutional guidelines determined by the Embryonic Stem Cell Research Oversight Committee (ESCRO) at Washington University in St. Louis. hESCs were routinely cultured in matrigel coated plates (Corning, Tewksbury, MA) in mTESR1 media (Stem Cell Technologies, Vancouver, Canada) supplemented with 1% pen strep (Gibco, Waltham, MA) and were kept in a humidified incubator at 37°C in 5% CO_2_ and 5% O_2_ levels. Wild-type and DC mutant hESCs were maintained and passaged onto new 6-well plates every 5 days at a split ratio of 1:12. The passage number of cells used in different experiments is indicated in each figure.

### Gene Editing

DKC1_A353V and DKC1_A353V_p53^−/−^, were generated using CRISPR/Cas9 genome editing technology and are described in detail in^9^. In brief, CRISPR gRNAs were inserted into the MLM3636 plasmid (Addgene 43860) and co-transfected with a plasmid carrying Cas9 (Addgene 43945) using the 4D-Nucleofector with the P4 Primary Cell 4D-Nucleofector kit (Lonza, Allendale, NJ). Single-stranded DNA donor oligos were co-transfected with plasmids carrying specific gRNAs and Cas9. For DKC1_A353V_p53^−/−^, one gRNA sequence was co-transfected with Cas9 to induce non-homologous end-joining (NHEJ) resulting in a frame shift and early termination, which was verified by targeted sequencing and protein expression analysis. Nucleofected cells were seeded on matrigel at low density and manually picked when colonies reached an appropriate size. Clones were then screened and sequenced. DKC1_A353V+TERC, and DKC1_A353V_shHNF4α hESCs were generated by zinc finger nuclease (ZFN). Transfection targeting the AAVS1 locus was performed with X-TremeGene 9 following the manufacturer’s instructions (Roche, Indianapolis, IN).

### Hepatocyte differentiation

Differentiation of hESCs into hepatocyte-like cells was performed according to^10^. Briefly, hESCs were transferred to a 100 mm plate coated with matrigel and incubated at 37°C in 5% CO_2_ and 5% O_2_ for 3 days or until 90-95% confluent. The cells were then transferred to 6 well plates previously covered with Matrigel and incubated with mTESR1. 24 hours later (Day 01) differentiation media [RPMI (Gibco), 1% Pen/Strep (Gibco), 1% non-essential amino acids (Gibco)] was added. On Day 02 of differentiation, 2% B27 minus insulin (Gibco), 100 ng/ml activin A (R&D Biosystems), 10 ng/ml BMP4 (R&D systems), and 20 ng/ml FGF (R&D Biosystems) was added to the media and cells were incubated in 20% O_2_. Between Day 03 and Day 05 of differentiation, only 2% B27 minus insulin and 100 ng/ml activin A were added to the differentiation media (changed daily). From Days 06 to 10, the differentiation media was supplemented with 2% B27 (Gibco), 20 ng/ml BMP4 and 10 ng/ml FGF. Between Days 06 and 10, cells were incubated at 5% O_2_. Between Days 11 and 15, cells were kept in differentiation media containing 2% B27 and 20 ng/ml HGF (Peprotech), again at 5% O_2_. From Day 16 forward, cells were kept in HCM medium (HCM Bullet Kit, Lonza) supplemented with “singlequots” supplied with the kit (except for the EGF) and 20 ng/ml Oncostatin-M (R&D Biosystems) at 20% O_2_. Cells and/or media were collected at days 06 (endoderm cells), 11 (hepatic endoderm), 16 (immature hepatocytes) and 21 (mature hepatocytes).

### Immunostaining

For immunostaining cells were fixed with 4% paraformaldehyde, permeabilized with 0.5% Triton X-100 and incubated with the following antibodies: OCT4 (Santa Cruz Biotechnology), TRA160 (Abcam), HNF4α (Abcam), AFP (R&D systems) and albumin-FITC (Dako). The cells were, then, incubated with appropriate secondary antibodies and nuclei were counterstained with DAPI. For quantification of albumin positive cells, we counted 1000 cells/slide on day 21.

### Quantification of albumin secretion

Albumin secretion quantification was performed using the Human Albumin ELISA kit from Bethyl Laboratories (Montgomery, TX), following manufacturers’ guidelines. Briefly, 100 µl of diluted (1:100) media from each sample was added to a 96-well plate. After 1-hour at room temperature, the plate was washed (1X “wash buffer”) and 100 µl of anti-albumin detection antibody was added to each well, following 1 h incubation at room temperature. After wash, 100 µl of “HRP Solution A” was added to each well. After a 30-minute incubation period at room temperature, wells were washed and “TMB Substrate Solution” was added. After a 30-minute incubation in the dark, the reaction was stopped by adding 100 µl of “Stop Solution” to each well. Absorbance was measured on a plate reader at 450 nm. Albumin concentration was quantified by interpolating their absorbance from the albumin standard curve generated with the samples.

### Immunoblots

Protein extraction was performed using NP-40 buffer (25 mM HEPES-KOH, 150 mM KCl, 1.5 mM MgCl2, 10% glycerol, 0.5% NP40, and 5 mM 2ME [pH 7.5]) supplemented with protease and phosphatase inhibitors (Roche, Indianapolis, IN) for 20 minutes on ice. Quantification of proteins was performed by Bradford assay. Proteins were resolved in 10% polyacrylamide gels in 1X Tris/glycine/SDS buffer and transferred onto nitrocellulose membrane at 400 milliamps for 1:45 hours in 1X Tris/glycine buffer with 20% methanol. Membranes were blocked in either 5% BSA or 5% milk in TBS buffer. Primary antibody incubation was performed overnight at 4°C in 5% BSA in TBS- buffer supplemented with 1% Tween-20 (TBS-T). Membranes were washed (3X 10 minutes) with TBS-T buffer and incubated in 1% milk in TBS-T with secondary antibodies (Li-COR, Lincoln, NE) for 1 hour. Membranes were then washed with TBS-T and scanned using the odyssey IR scanner (Li-COR). Image capture and signal analysis was done using the Image Studio software (Li-COR). Primary antibodies used in this study were: phospho-H2AX (1:1000, Abcam, Cambridge, MA), p53 (1:1000, Santa Cruz Biotechnology, Santa Cruz, CA) and Actin (1:2000, Sigma, St. Louis, MO).

### Detection of Telomerase Activity

Telomerase activity was measured by Telomere Repeat Amplification Protocol (TRAP). Briefly, cells were lysed in NP-40 buffer for 20 min on ice and extracts clarified by centrifugation at 16,000g for 10 min. Protein quantification was performed by Bradford assay. Telomere extension reactions were performed using 2.0 μg, 0.5 μg and 0.125 μg of protein and resulting products were amplified by PCR, following a modified 2-step TRAP protocol from the manufacturer (TRAPeze, EMD Millipore, Boston, MA).

### Telomere length analysis

Telomere length was quantified by Telomere Repeat Fragment Analysis (TRF). Isopropanol-extracted DNA was digested (10µg) overnight with RSA and HINF1 restriction enzymes (New England Biolabs, Ispwich, MA) and resolved (2.5µg for each analysis) on a 0.8% agarose gel for 16 hours at 85 volts in TBE (Tris/Borate/EDTA) buffer. The gel was then soaked in denaturing buffer (1.5M NaCl and 0.5M NaOH) for 45 minutes followed by neutralizing buffer (1.5M NaCl, 1M Tris-HCL at pH 7.4) for 1 hour. DNA was transferred to a nitrocellulose membrane by capillarity for at least 16 hs in 20x saline-*sodium citrate* (3M NaCl, 0.3M sodium citrate dehydrate at pH 7.0). After cross-linking, the membrane was hybridized with a ^32^P-labelled probe (TTAAGGG)_4_ and exposed overnight to Carestream BioMax MR film (Sigma).

### Flow Cytometry

Flow cytometry analysis was done on BD LSR Fortessa at the Department of Pathology & Immunology Flow Cytometry Core (Washington University in St. Louis). Antibodies used were the following: CXCR4-PE (BD Biosciences), CD117-APC (Invitrogen).

### RNA extraction, cDNA synthesis, and quantitative real time PCR analysis

RNA extraction was performed using Trizol (Invitrogen) following manufacturer’s instructions. cDNA synthesis was made using Superscript III First Strand synthesis kit (Invitrogen) following manufacturer’s instructions. Quantitative real-time PCR was performed using a StepOne Plus (Applied Biosystems, Waltham, MA) instrument. For transcriptional analysis of all coding genes, Evagreen master mix (Lambda Biotech, St Louis, MO) was used and reactions were performed in duplicates with 100ng of cDNA per reaction. For TERC qRT-PCR analysis, Brilliant II 1-step qPCR master mix (Agilent, Santa Clara, CA) was used following manufacturer’s instructions. Reactions were performed in duplicates with 100ng of RNA per reaction. Sample size for all experiments was at least *n*=3. Expression levels were calculated by ΔΔCT. β-actin was used as the reference loading gene. All primer sequences can be found in (Supplemental Table 1).

### Caspase Activity Measurement

Caspase activity was quantified using dedicated Caspase 3, 8, and 9 colorimetric detection kits following manufacturer’s instructions (Abcam).

## RESULTS

### Telomerase is quickly down-regulated during hepatocyte differentiation from hESCS

Our *in vitro* hepatic-derivation protocol from hESCs (adapted from^11^) recapitulates different hepatocyte development stages including endoderm (day 6; *CXCR4, SOX17* positive cells), hepatic endoderm (day 11; *HNF4α* positive cells), immature hepatocytes (day 16; *AFP* positive cells) and finally mature, hepatocyte-like cells (day 21) that express *FGA, FGG, CYP1A1* and albumin (Figure 1A-B). Recapitulating *in vivo* data, telomerase activity is quickly down-regulated during the initial stages of differentiation (Figure 1C), due to the specific silencing of *TERT* expression, as *TERC* levels remain unaltered (Figure 1D). Importantly, despite the fast silencing of telomerase, telomeres are not significantly shortened during the 21 days of hepatic differentiation (Figure 1E), indicating that the initial telomere length at Day 1 of differentiation can be used as a reference for observed phenotypes during the entire, 21 days, hepatocyte maturation protocol.

**Figure 1:**
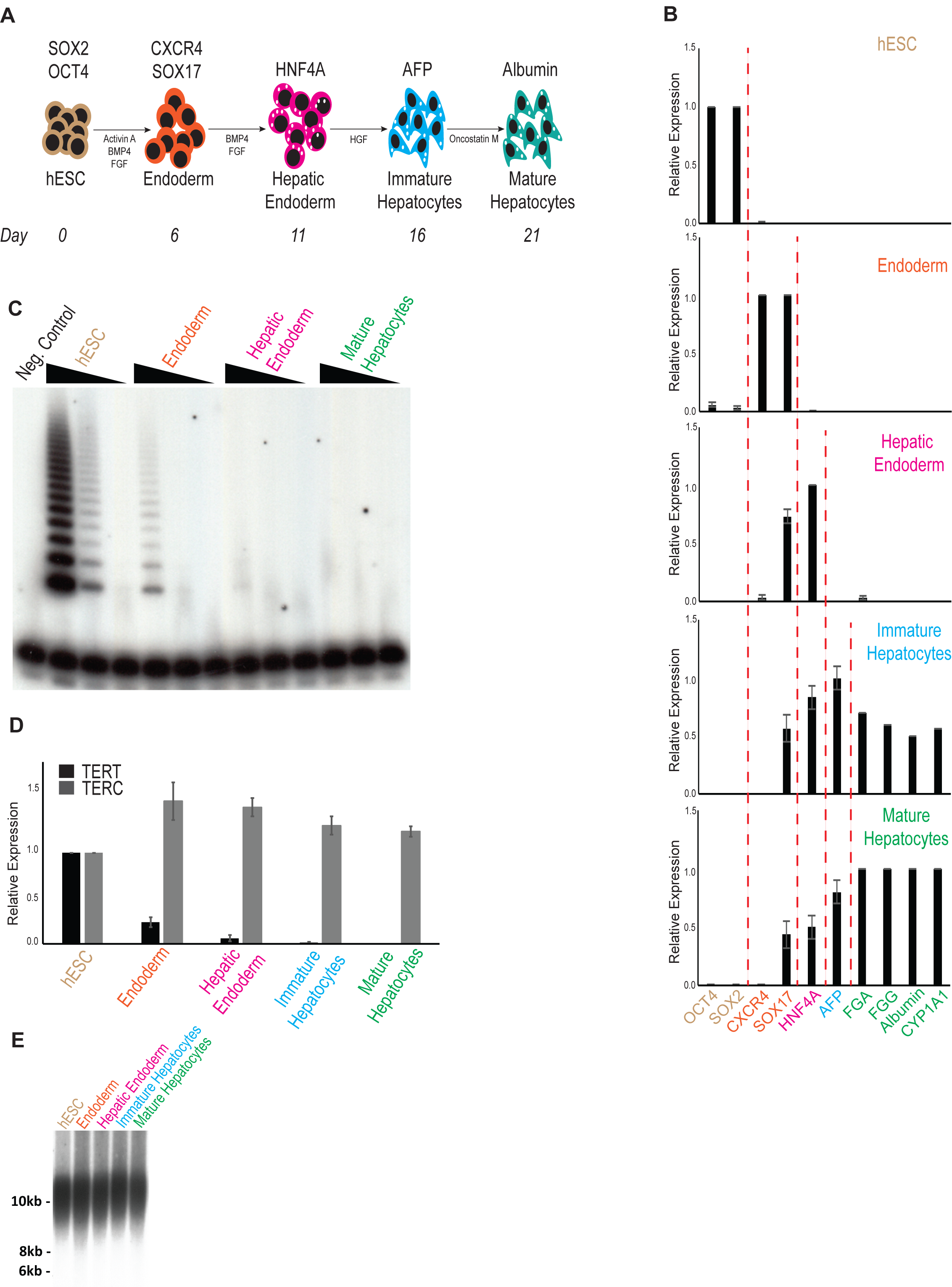
Telomerase is quickly down regulated during hepatocyte differentiation of hESCs. **(A)** Schematic of *in vitro* hepatocyte differentiation. Hepatocyte differentiation is achieved through serial addition of specific cytokines and growth factors, depicted in the model. During hepatic differentiation, different transient cell populations can be identified and isolated at specific times (indicated in the bottom line). These include Endoderm, Hepatic Endoderm, Immature Hepatocytes and Mature Hepatocytes. Hepatocyte differentiation protocol adapted from^10^. **(B)** Relative gene expression analysis by real-time quantitative PCR of different markers specific for each cellular population obtained during hepatic differentiation: *OCT4* and *SOX2*, hESCs (Day 0); *CXCR4* and *SOX17*, Endoderm (Day 6); *HNF4α*, Hepatic Endoderm (Day 11); *AFP*, Immature Hepatocytes (Day 16); *FGA, FGG*, Albumin and *CYP1A1*, Mature Hepatocytes (Day 21). **(C)** Telomerase activity by TRAP during hepatic differentiation from WT hESCs. Range of concentrations represent four-fold serial dilutions. Negative control: NP40 buffer. **(D)** Quantitative Real-Time PCR analysis of the telomerase core components *TERT* and *TERC* during hepatic differentiation from WT hESCs. **(E)** Telomere length analysis by Telomere Restriction Fragment (TRF) during hepatic differentiation from WT hESCs. Molecular weight (in kb) is shown.

### Progressive telomere shortening impairs hepatocyte development

Combining our genetically engineered DKC1_A353V hESCs with this *in vitro* hepatocyte differentiation protocol, we were able to study hepatocyte development in clinically relevant telomerase mutant cells with progressively shorter telomeres. We chose two different stages, DKC1_A353V cells with longer telomeres (referred to as early passage –EP; passage <13) and shorter telomeres (referred to as late passage – LP; passage >30; Supplemental Figure 1G) to understand the consequences of progressive telomere shortening in hepatocyte differentiation and function. Telomere length did not interfere with initial endoderm formation, measured both by quantification of CXCR4/CD117+ cells by flow cytometry (Figure 2A) and by the expression of *FOXA2* and *SOX17* by quantitative real-time PCR (Figure 2B). However, telomere dysfunction severely compromised formation of hepatic endoderm, as *HNF4α* levels were significantly reduced in DKC1_A353V_LP cells when compared to WT and DKC1_A353V_EP cells (Figure 2C). Accordingly, the generation of mature, hepatocyte-like cells from DKC1_A353V in late passage was significantly reduced when compared to WT and DKC1_A353V_EP cells, with low expression levels of hepatocyte markers (Figure 2D), albumin positive cells (Figure 2E-F), and albumin secretion (Figure 2G) on day 21 of differentiation.

**Figure 2:**
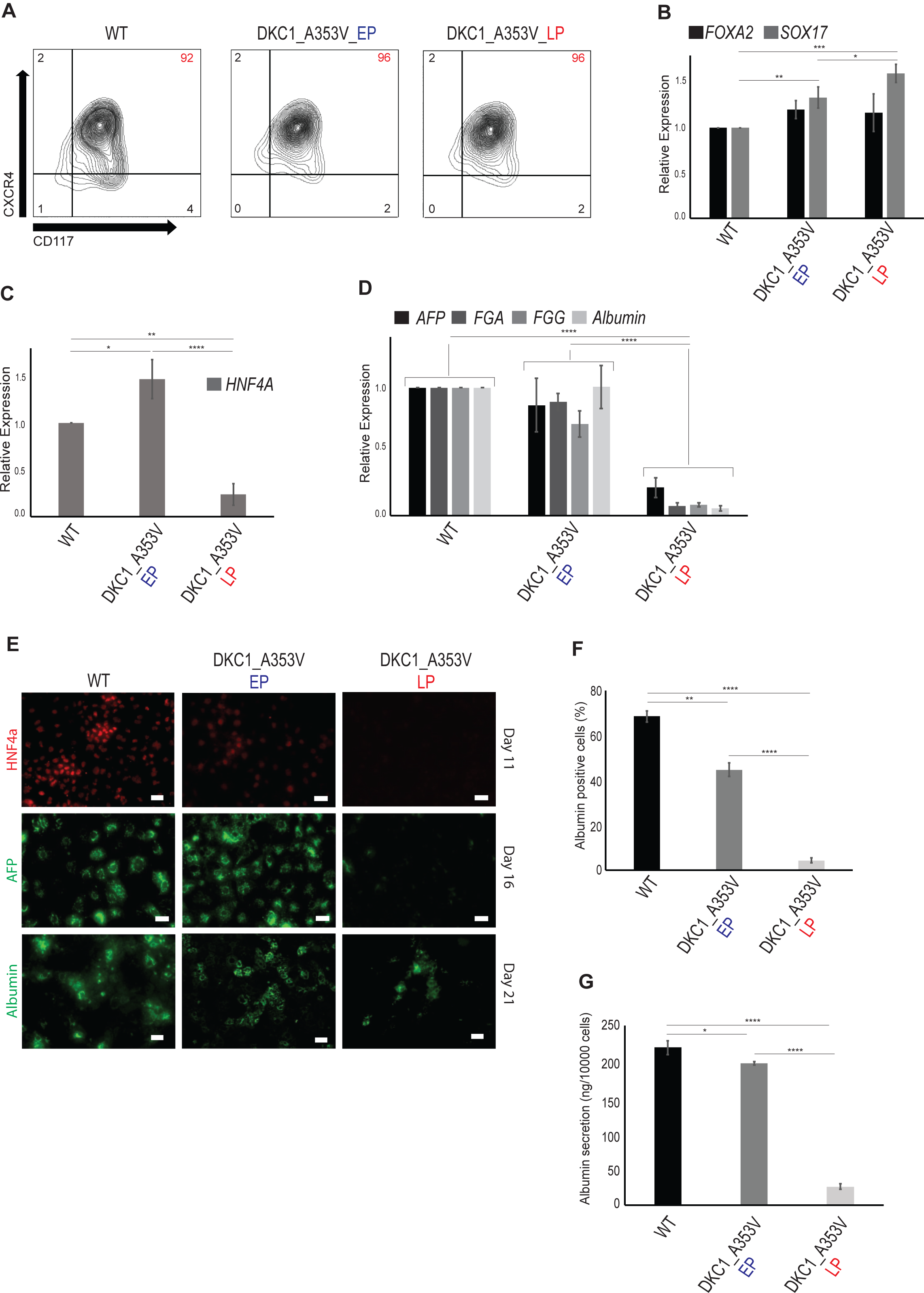
Telomere shortening impairs hepatocyte differentiation in DKC1_A353V hESCS. **(A, B)** Endoderm generation from WT and DKC1_A353V hESCs at early (EP) and late (LP) passages, as assessed by **(A)** formation of CXCR4+CD117+ population by flow cytometry (% of population of interest indicated in red, top right) and **(B)** relative expression (analyzed by quantitative RT-PCR) of the endoderm markers *FOXA2* and *SOX17.* **(C)** Generation of an hepatic endoderm population quantified by the relative gene expression of *HNF4α* (by quantitative RT-PCR) in WT and DKC1_A353V cells at different passages. **(D)** Relative expression of hepatocyte markers (by quantitative RT-PCR) after 21 days of differentiation in WT and DKC1_A353V cells at different passages. **(E)** Immunocytochemistry showing expression of different markers (indicated in the figure) during differentiation of WT and DKC1_A353V cells in early or late passages. The specific day during differentiation is indicated on the right. Scale Bars represent 50 μM. **(F)** Quantification of albumin positive cells by immunofluorescence after 21 days of differentiation in WT and DKC1_A353V cells at different passages. A total of 1000 cells/slide was counted. **(G)** Quantification of albumin secretion (by ELISA) after 21 days of differentiation in WT and DKC1_A353V cells at different passages. *n*=3, mean ± SEM, *p≤0.05; **p≤0.0025; ****p≤0.0001. Statistical analysis was performed using one-way ANOVA followed by Tukey’s post hoc test.

In addition, we wanted to understand if dyskerin’s well established role in ribosomal biology^12^ could also be responsible for the reduced hepatocyte differentiation efficiency observed in DKC1_A353V cells. We therefore restored telomere length in DKC1_A353V_LP cells, by expressing *TERC* from the safe harbor AAVS1 locus^13^ (Supplemental Figure 2A-B). It is clear that restoring telomere maintenance in cells that retain mutant dyskerin efficiently rescues hepatic endoderm formation, as well as hepatocyte derivation (Supplemental Figure 2C-F), therefore establishing telomere shortening, and not impaired ribosomal biology, as the causative effect for the reduced hepatocyte differentiation observed in cells harboring mutations in dyskerin.

### Stabilization of p53 reduces hepatocyte differentiation from DKC1_A353V hESCs

As telomere shortening is a known inducer of senescence and cell death, we next decided to understand if the failure to form hepatic endoderm in our DKC1_A353V hESCs with short telomeres was a consequence of reduced cellular viability. Indeed, we verified that during hepatocyte differentiation of DKC1_A353V_LP hESCs, there is accrual of DNA damage (as measured by γH2AX accumulation) and stabilization of p53 (Figure 3A), both of these common responses of telomere shortening and dysfunction in mammalian cells. Surprisingly however, our data indicates that the reduced hepatocyte specification observed in DKC1_A353V_LP cells was not caused by apoptosis activation (Figure 3B) or reduced cellular proliferation (Figure 3C). In fact, while DKC1_A353V_LP are unable to differentiate into hepatocyte-like cells (Figure 2D-G), overall cellular proliferation was significantly increased in DKC1_A353V_LP cells during the 21 days of differentiation (Figure 3C). This indicates that telomere shortening specifically impairs hepatocyte development through a mechanism that is independent from its well established role in cell death and/or cell cycle arrest.

**Figure 3:**
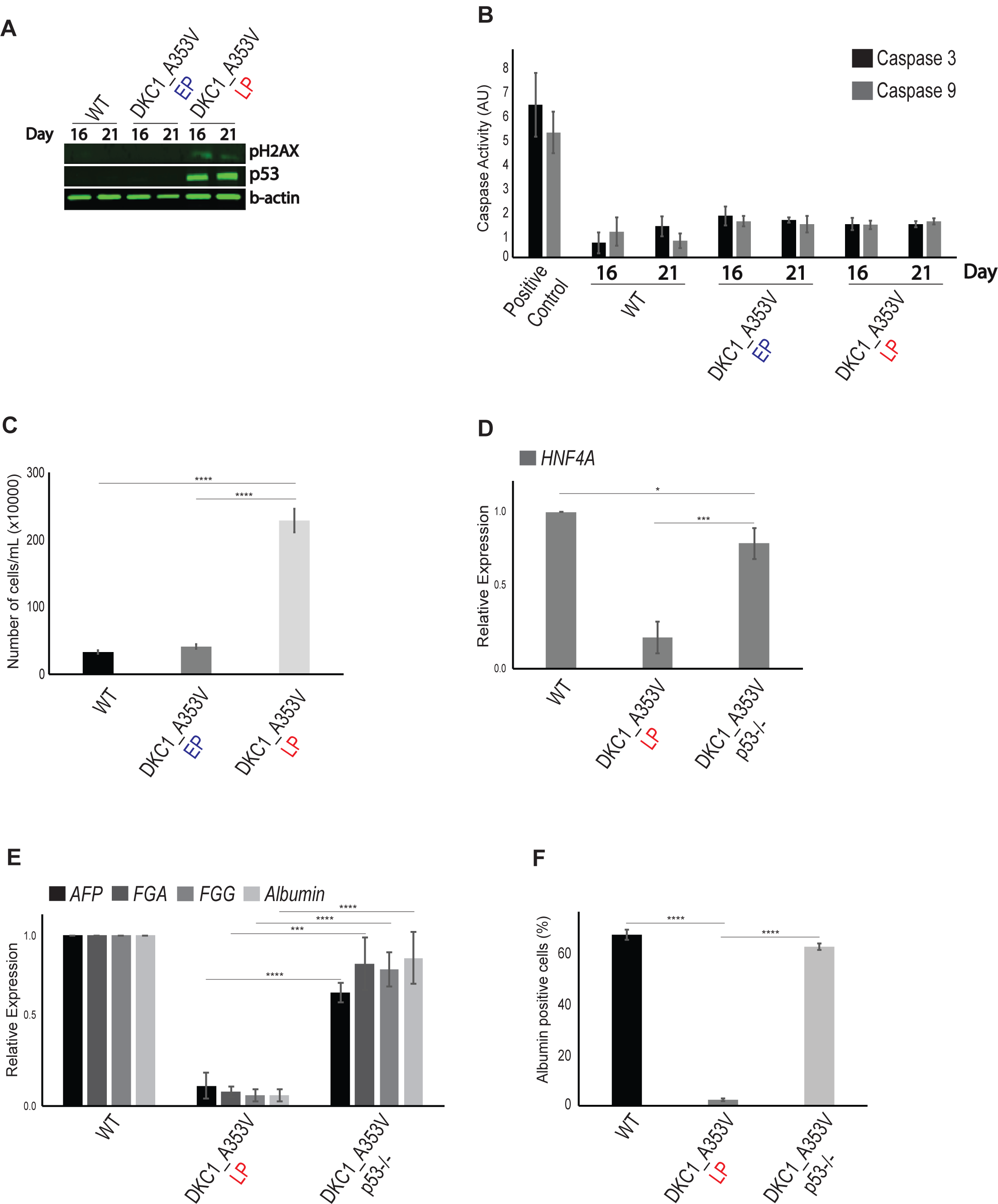
p53 stabilization significantly reduces hepatocyte derivation from DKC1_A353V mutant hESCs. **(A)** Representative immunoblot analysis of γH2AX and p53 expression on days 16 and 21 of hepatic differentiation (immature and mature hepatocyte stages) of WT and DKC1_A353V hESCs (at early and late passages). β-actin is shown as loading control. **(B)** Quantification of caspases 3 and 9 activation in early passage and late passage cells after 21 days of differentiation. Positive control: Ultraviolet Light irradiated (UV; 30J/m^2^) WT hESCs. **(C)** Total number of cells after hepatic differentiation of WT and DKC1_A353V hESCs. Cells were collected on day 21 and figure shows total number of cells found in each population (total numbers quantified by cell counter). **(D)** Generation of hepatic endoderm population quantified by the relative gene expression (by quantitative RT-PCR) of *HNF4α* in WT, DKC1_A353V_LP and the p53 ablated DKC1_A353V_p53^−/−^ cells. **(E)** Relative expression of hepatocyte markers by qRT-PCR after 21 days of differentiation (mature hepatocyte stage) in WT, DKC1_A353V_LP and DKC1_A353V_p53^−/−^ cells. **(F)** Quantification of albumin positive cells by immunofluorescence after 21 days of differentiation in WT, DKC1_A353V_LP and DKC1_A353V_p53^−/−^ cells. *n*=3, mean ± SEM, *p≤0.05; **p≤0.0025; ***p≤0.001; ****p≤0.0001. Statistical analysis was performed using one-way ANOVA followed by Tukey’s post hoc test.

We therefore decided to probe if the observed p53 up-regulation during the hepatocyte differentiation of DKC1_A353V_LP hESCs cells (Figure 3A) was a determinant factor in the reduced efficiency of hepatocyte development observed in cells with short telomeres. For that we used DKC1_A353V_LP hESCs where we ablated p53 through genome engineering (CRISPR/cas9)^10^. These DKC1_A353V_p53^−/−^ hESCs retain short telomeres, as telomerase is still defective, but have no activation of p53-dependent signaling^10^. Interestingly, although retaining short telomeres, these cells have normal hepatocyte differentiation capacity, with restored levels of *HNF4α* expression during the hepatic endoderm stage (Figure 3D) and restored expression of hepatocyte markers (Figure 3E). Likewise, there is a significant increase in the number of albumin positive cells at the mature hepatocyte-like cells stage (Figure 3F). Collectively, these data indicate that the activation of *p53* by telomere shortening specifically impairs hepatocyte development from hESCs.

### Restoring HNF4α successfully rescues hepatocyte derivation and function in telomerase mutant cells

As the earliest phenotype observed during the hepatic differentiation of DKC1_A353V hESCs with short telomeres was reduced hepatic endoderm formation, we focused more deeply on that developmental stage. Specifically, we hypothesized that the marked reduction in *HNF4α* expression in DKC1_A353V cells could be caused by p53 stabilization, as it has been previously suggested in transformed cells^14^. HNF4α is a key regulator of hepatocyte differentiation during embryonic development and maintenance of a differentiated phenotype in adult livers, as it regulates over 60% of genes involved in hepatocytes^15^. Disruption of HNF4α in mature hepatocytes is linked to epithelial to mesenchymal transition (EMT)^16, 17^ and increased cellular proliferation^15, 18^.

In our *in vitro* differentiation system we verified that *HNF4α* expression in WT cells increases continuously until day 11 of differentiation (Figure 4A). Moreover, the silencing *HNF4α* in WT cells by constitutive expression of short-hairpin RNA sequences from the AAVS1 locus (Supplemental Figure 3), causes a significant reduction in hepatic differentiation (Figure 4B), similar to what we observe in DKC1_A353V_LP cells. Therefore, to confirm that the reduced expression of *HNF4α* was responsible for the reduced hepatic differentiation observed in DKC1_A353V_LP cells, we again used genome engineering to conditionally express (in a DOX-inducible system; Figure 4C, inlet) *HNF4α* in late passage DKC1_A353V cells. Surprisingly, the expression of *HNF4α* in cells that retain short telomeres (Figure 4C) is able to rescue hepatic differentiation (Figure 4D) and function (Figure 4E). Moreover, expression of *HNF4α* prevents the increased cellular proliferation observed in cells with dysfunctional telomeres (Figure 4F). Combined, this data indicates that restoring *HNF4α* levels successfully overcomes the deleterious consequences of telomere dysfunction observed during hepatocyte development.

**Figure 4:**
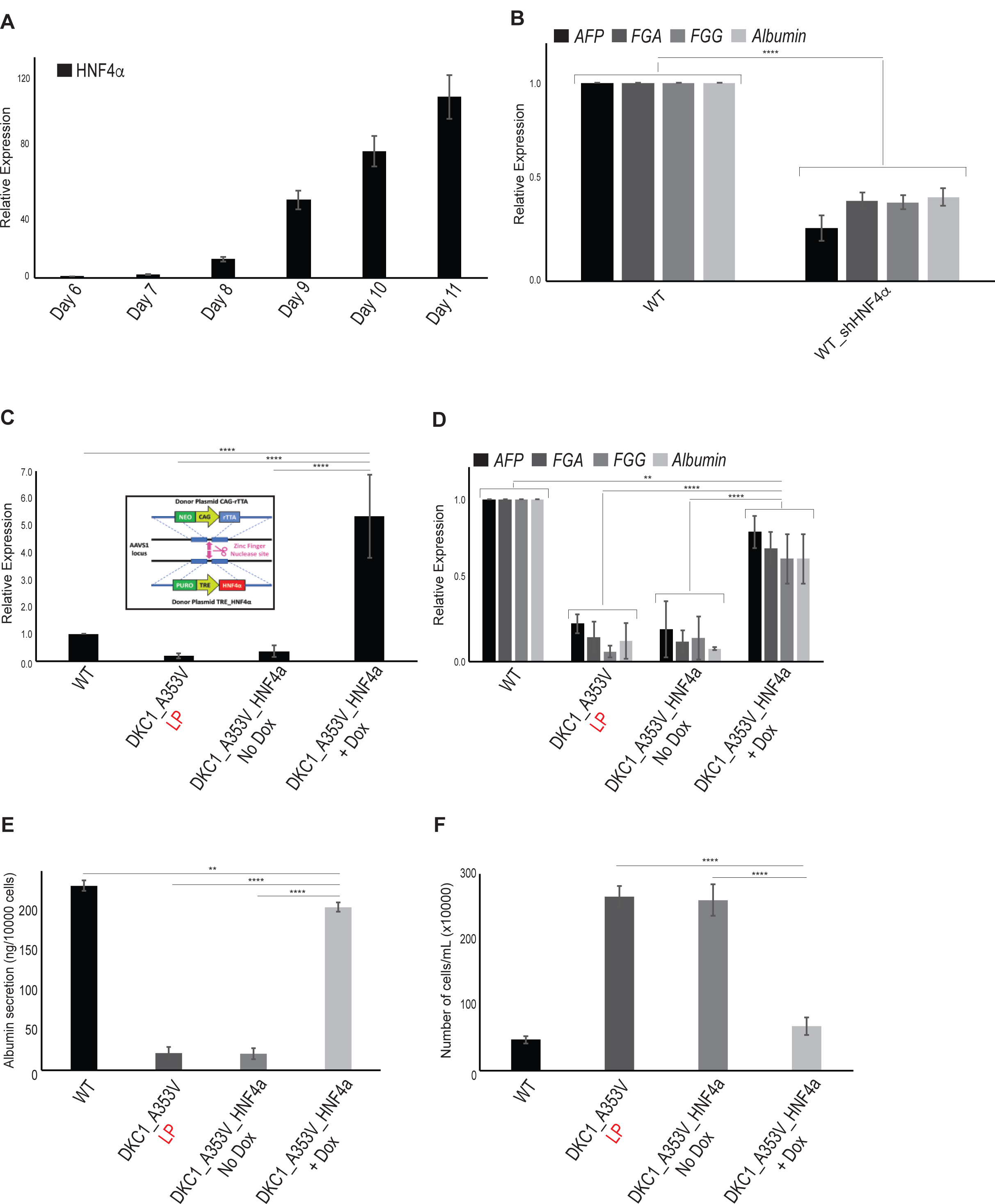
HNF4a expression rescues hepatocyte development in DKC1_A353V mutant cells that retain short telomeres. **(A)** *HNF4α* expression during hepatocyte derivation from WT hESCs. *HNF4α* expression increases until day 11 of differentiation, at the hepatic endoderm stage. **(B)** Relative expression of hepatocyte markers by qRT-PCR after 21 days of differentiation (mature hepatocyte stage) in WT and WT_sh HNF4α cells. **(C)** *HNF4α* expression in WT, DKC1_A353V_LP, and conditional DKC1_A353V_ *HNF4α* cells with or without addition of Doxycycline. *Inlet:* construction and cloning of conditional HNF4α cassette into the AAVS1 locus of DKC1_A353V hESCs (model adapted from^12^). **(D)** Relative expression (by quantitative RT-PCR) of hepatocyte markers after 21 days of differentiation in WT, DKC1_A353V_LP, and conditional DKC1_A353V_ *HNF4α* cells with or without addition of Doxycycline. **(E)** Quantification of albumin secretion after 21 days of differentiation in WT, DKC1_A353V_LP, and conditional DKC1_A353V_ *HNF4α* cells with or without addition of Doxycycline. **(F)** Total number of cells after hepatic differentiation of WT, DKC1_A353V_LP, and conditional DKC1_A353V_ *HNF4α* cells with or without addition of Doxycycline. Cells were collected on day 21 and figure shows total number of cells found in each population (total numbers quantified by cell counter). *n*=3, mean ± SEM, *p≤0.05; **p≤0.0025; ***p≤0.001; ****p≤0.0001. Statistical analysis was performed using one-way ANOVA followed by Tukey’s post hoc test.

## CONCLUSIONS

Combined, our data clearly establishes *HNF4α* levels as a major determinant of hepatic differentiation in settings of short telomeres and provides insight on the mechanism behind the severe hepatocyte failure observed in patients with telomere syndromes. As low levels of *HNF4α* are commonly found in liver disease, our data indicates that *p53* mediated silencing of *HNF4α* could be linked to disease onset in patients with defective telomere maintenance and, therefore, potential therapies aiming at restoring *HNF4α* levels in these patients could prove beneficial. In this regard, while further analysis is necessary to decipher the specific mechanism of HNF4α silencing by *p53*, our *in vitro* differentiation platform provides a robust platform for genetic and pharmacologic screening for hepatocyte development and function in settings of damaged telomeres.

## AUTHORSHIP CONTRIBUTIONS

E.L.N., W.C.F., and L.F.Z.B. designed the experiments and analyzed the data; E.L.N., A.T.V., W.C.F., K.A.B. and L.F.Z.B. performed the experiments; E.L.N. and L.F.Z.B. wrote the manuscript.

## CONFLICT OF INTEREST

Nothing to disclose

## ACKNOWLEDGEMENTS

E.L.N. was supported by CNPq, Brazil. A.T.V was supported by the Phillip Majerus Postdoctoral Fellowship. W.C.F. was supported by NHLBI T32 Training Grant in Molecular Hematology (HL007088-41). K.A.B. was supported by the National Fellowship Foundation. L.F.Z.B. is supported by the NHLBI (R01HL137793-01), the Department of Defense (DOD; BM160054) and grants from the V Foundation for Cancer Research, Edward Mallinckrodt Jr. Foundation, Concern Foundation, American Federation for Aging Research (AFAR) and the Longer Life Foundation. This project was also supported by a pilot grant from the Washington University DDRCC program (NIDDK P30 DK052574) to L.F.Z.B.

