## Supplementary figures and images for "Telomere dysfunction represses HNF4α leading to impaired hepatocyte development and function"

### Supplemental Figures

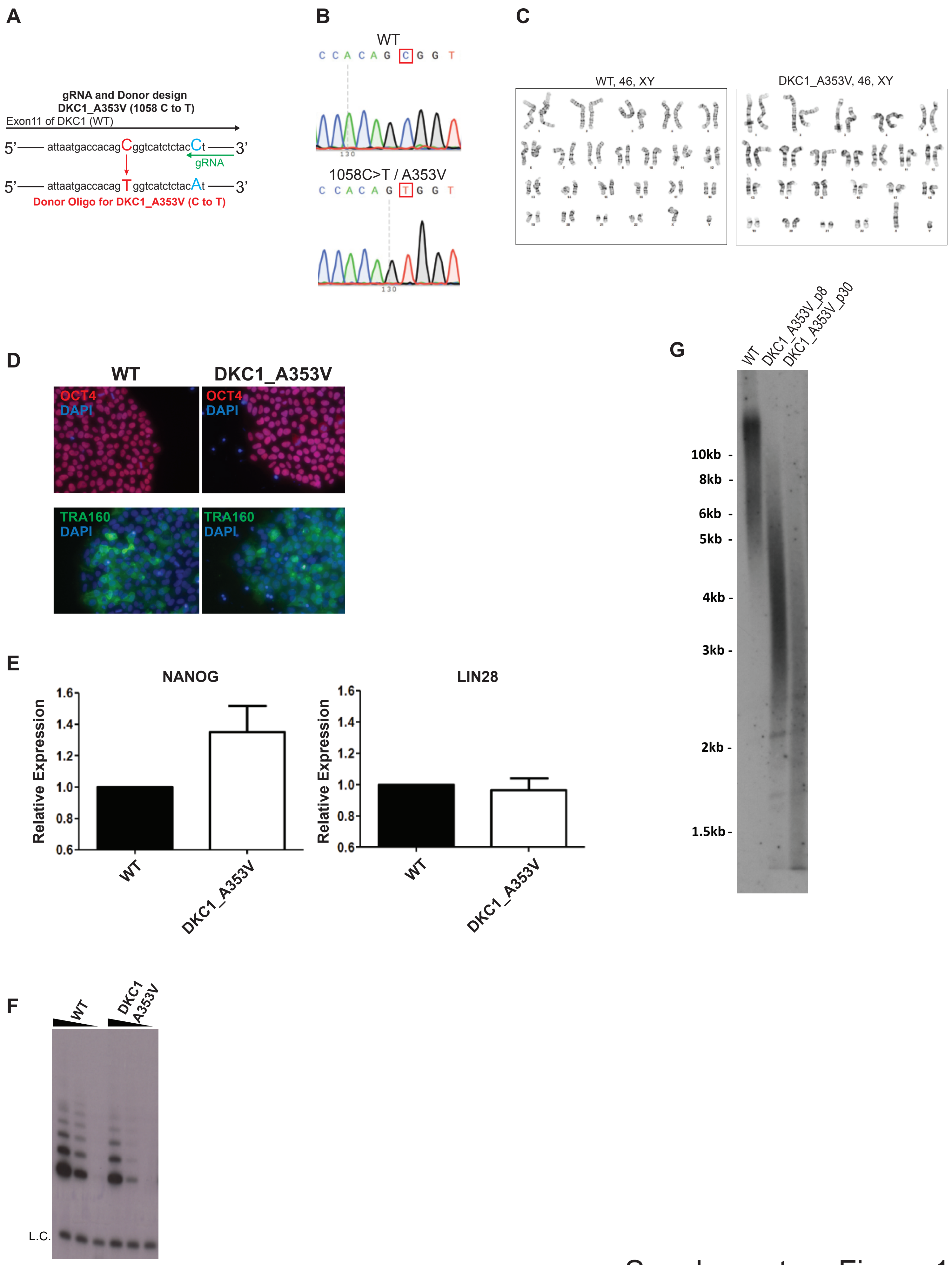


Supplementary Figure 1

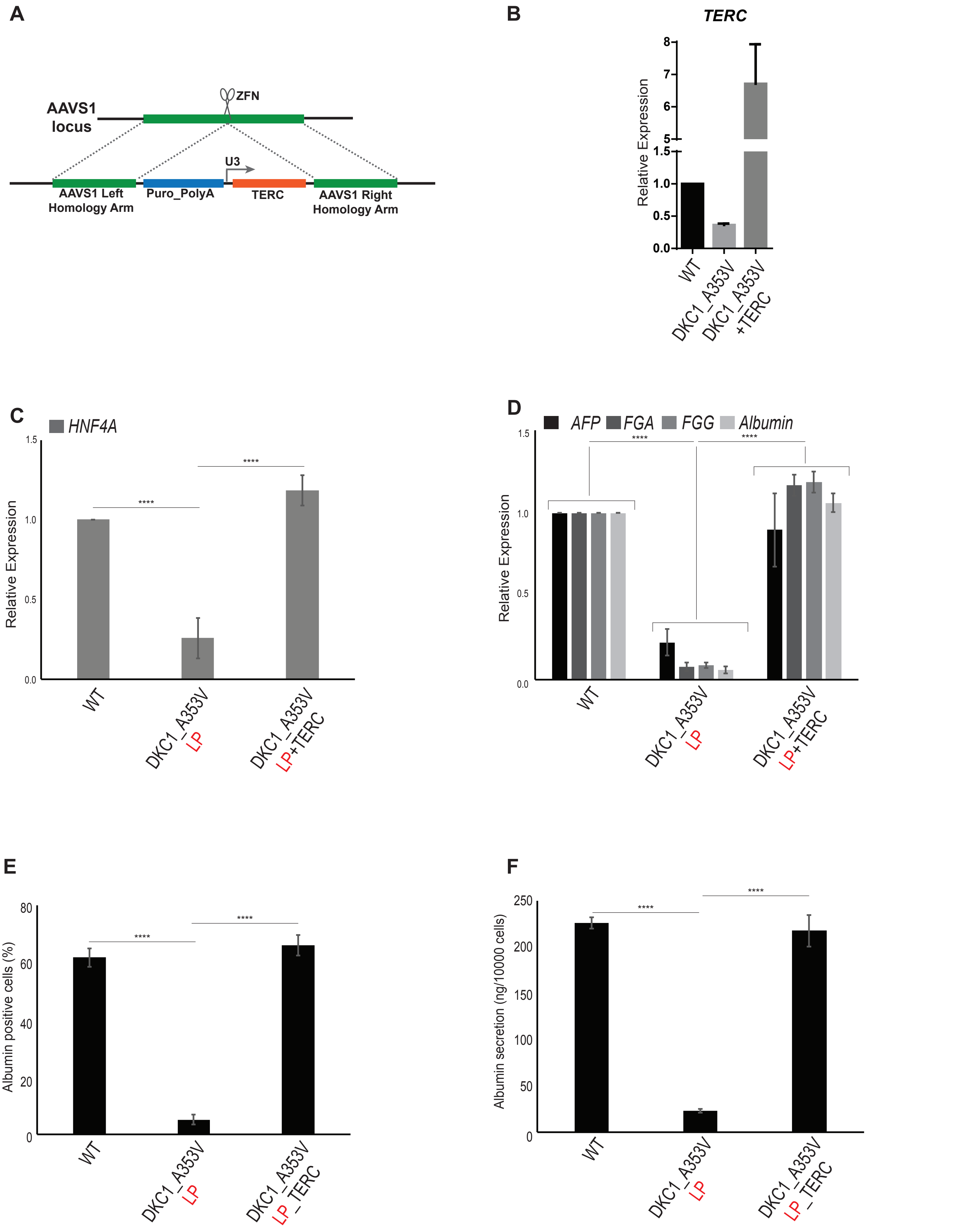


Supplementary Figure 2

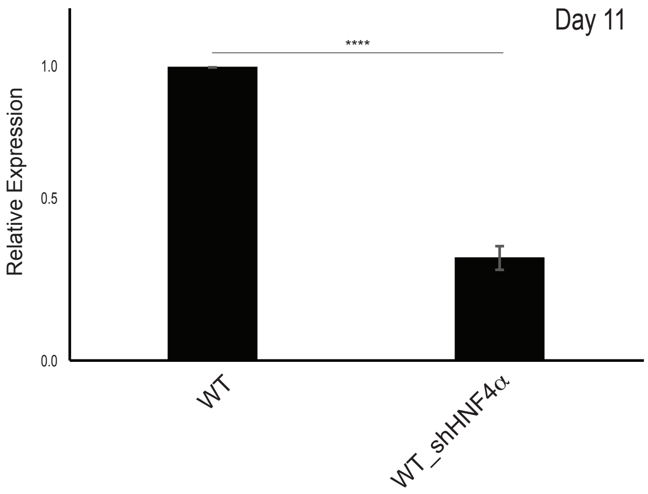


Supplementary Figure 3
