## Supplemental Text for "Telomere dysfunction represses HNF4α leading to impaired hepatocyte development and function"

**SUPPLEMENTARY FIGURE LEGENDS**

**Supplementary Figure 1: Construction of telomerase mutant DKC1_A353V hESCs.** **(A)** Strategy for introduction of disease-specific mutations in DKC1. Guide RNAs (gRNAs) targeting Exon 11 (DKC1) were used with in combination with specific ssDNA donor oligo templates for introduction of DKC1 (A353V; *C>T*). In blue, silent mutations introduced to facilitate CRISPR/Cas9 mediated genome modification. A detailed description of engineering of DKC1_A353V hESCs can be found in^9^. **(B)** DNA sequencing confirms correct C>T modification in DKC1_A353V hESCs. **(C)** G-band analysis in wild-type and DKC1_A353V hESCs. No chromosomal abnormalities were detected. **(D)** Immunofluorescence analysis confirming normal expression of the pluripotency markers OCT4 and TRA160 in DKC1_A353V hESCs. **(E)** Quantitative Real-Time PCR analysis of different pluripotency markers in WT and DKC1_A353V hESCs. **(F)** Telomerase activity by TRAP in WT and DKC1_A353V mutants. Range of concentrations represent four-fold serial dilutions. L.C: loading control. (G) Telomere length analysis by Telomere Restriction Fragment (TRF) of wild-type and DKC1_A353V hESCs at different cell passages, demonstrating progressive telomere shortening in mutant cells. Molecular weight (in kb) is shown.

**Supplementary Figure 2: Expression of *TERC* rescues hepatocyte derivation from DKC1_A353V hESCs.** (**A)** Model of AAVS1 targeting in DKC1_A353V hESCs. *TERC* is expressed under the control of an U3 promoter sequence in DKC1_A353V+TERC hESCs. For more details on this construct, please see reference^9^. **(B)** Quantification of *TERC* levels by qRT-PCR in wild-type, DKC1_A353V, and DKC1_A353V+TERC hESCs. **(C)** Quantification of *HNF4α* levels by qRT-PCR on Day 11 of differentiation (Hepatic Endoderm stage) of wild-type, DKC1_A353V, and DKC1_A353V+TERC cells. **(D)** Relative expression of hepatocyte markers by qRT-PCR after 21 days of differentiation (mature hepatocyte stage) in WT, DKC1_A353V_LP and DKC1_A353V+TERC cells. **(E)** Quantification of albumin positive cells by immunofluorescence after 21 days of differentiation in WT, DKC1_A353V_LP and DKC1_A353V+TERC cells. **(F)** Quantification of albumin secretion by ELISA reading after 21 days of differentiation in WT, DKC1_A353V_LP and DKC1_A353V+TERC cells. *n*=3, mean ± SEM, ****p≤0.0001. Statistical analysis was performed using one-way ANOVA followed by Tukey’s post hoc test.

**Supplementary Figure 3: Generation oh WT_shHNF4α hESCs.** Efficient silencing of *HNF4α* expression in WT_sh HNF4α cells at the hepatic endoderm stage (Day 11).
