## Supplemental table for "Telomere dysfunction represses HNF4α leading to impaired hepatocyte development and function"

| **mRNA** | **Forward sequence** | **Reverse sequence** |
| --- | --- | --- |
| FOXA2 | GGAGCAGCTACTATGCAGAGC | CGTGTTCATGCCGTTCATCC |
| SOX17 | CTCCGGTGTGAATCTCCCC | CACGTCAGGATAGTTGCAGTAAT |
| HNF4A | ATAGCTTGACCTTCGAGTGC | TGGACAAAGACAAGAGGAACC |
| AFP | GGCAGCCACAGCAGCCACTT | TGCAGCGCTACACCCTGAGC |
| FGA | CAGCCCCACCCTTAGAAAAG | CTCCTTCAGCTAGAAAGTCACC |
| FGG | CAAAGACACGGTGCAAATCC | TTCCAGACCCATCGATTTCAC |
| Albumin | TGGCACAATGAAGTGGGTAA | CTGAGCAAAGGCAATCAACA |
| TERC | CGCTGTTTTTCTCGCTGACT | GCTCTAGAATGAACGGTGGAA |
| TERT | CGAAAACCTTCCTCAGGACCC | GGCCGGCATCTGAACAAAAG |

**Sup. Table 1: Primer sequences used in this manuscript.**
